# Glycoprotein C Mutations Regulate the Plaque Size of Fusion-Defective Herpes Simplex Virus

**DOI:** 10.64898/2026.01.26.701780

**Authors:** Qing Fan, Richard Longnecker, Sarah A. Connolly

## Abstract

Herpes simplex virus 1 (HSV-1) entry and virus-induced cell fusion require the coordinated activity of the essential glycoproteins gD, gH/gL, and gB. Mutations in this core fusion machinery reduce viral entry, fusion, and spread. Previously, we showed that introducing three mutations in gB (gB3A) or replacing HSV-1 gB with saimiriine herpesvirus 1 (SaHV-1) gB resulted in fusion-defective, small-plaque viruses. Serial passage of these viruses selected mutations in gH and gC, and we demonstrated that the gH mutations partially restored fusion function. In this study, we examined whether mutations in gC contribute to plaque formation, fusion, and entry. We generated an additional fusion-defective, small-plaque gD chimeric virus by replacing the profusion domain of HSV-1 gD with the corresponding region from SaHV-1 gD (gDchi). Serial passage of this virus partially restored plaque size and also selected for a mutation in gC. This study demonstrates that the selected gC mutants contributed to the partial restoration of plaque size. Plaque size of all three fusion-defective viruses was enhanced by exogenous expression of the corresponding selected gC mutants, whereas exogenous expression of wild-type gC reduced plaque size of the passaged isolates carrying gC mutations. Although gC mutants did not enhance fusion or entry mediated by the wild-type core fusion machinery, both wild-type and mutant gC enhanced cell-cell fusion when coexpressed with fusion-defective gH mutants. These results indicate that gC functions as a conditional accessory regulator that can partially compensate for defects in the HSV-1 core fusion machinery.

**IMPORTANCE:** Herpes simplex virus 1 (HSV-1) entry is mediated by a core fusion machinery comprised of glycoproteins gD, gH/gL, and gB. The contributions of additional viral glycoproteins to this process are not well defined. Glycoprotein C (gC) is known to function in cell attachment to heparan sulfate and immune evasion but is dispensable for viral entry under standard conditions. Here, we show that gC can act as a conditional accessory regulator of HSV-1 entry and fusion. Using serial passage of fusion-defective viruses, we demonstrate that mutations in gC are repeatedly selected and can partially compensate for defects in entry and fusion. These findings indicate that gC contributes to viral fusion under suboptimal conditions and highlight a role for this accessory glycoprotein in the adaptability of herpesvirus entry.

## INTRODUCTION

Herpes simplex virus type 1 (HSV-1) is an alphaherpesvirus that causes recurrent mucocutaneous lesions and, less frequently, severe diseases including encephalitis and meningitis. Entry of HSV-1 into host cells and virus-induced cell-cell fusion require coordinated interactions among four essential glycoproteins, glycoprotein D (gD), the heterodimer gH/gL, and the fusion protein gB (1–3). The current model of HSV entry proposes that following attachment, binding of gD to a cellular receptor triggers conformational changes in gD that signal gH/gL, which in turn activates gB to drive membrane merger (4). All herpesviruses share the conserved entry glycoproteins gH/gL and gB, with receptor binding carried out by virus-specific glycoproteins.

We previously used a genetic approach to define functional interactions among these HSV-1 entry glycoproteins by exchanging homologous entry glycoproteins between HSV-1 and saimiriine herpesvirus 1 (SaHV-1) (5–7). These cross-species swap experiments revealed a species-specific functional interaction between gD and gH/gL, whereas gB from either virus was compatible with the entry machinery of both viruses. By generating chimeric glycoproteins composed of HSV-1 and SaHV-1 sequences, we identified species-specific functional interaction sites within the N-terminal domains of gH and the “profusion domain” of gD (5, 6).

Although gD, gH/gL, and gB are sufficient for fusion, additional viral proteins can modulate fusion efficiency (8–12). For example, mutations in gK, UL20, or UL24 can produce a syncytial plaque phenotype (13, 14). gK interacts with gB(15), and deletion of the gK or its N-terminus disrupts entry and virus-induced cell fusion (16–19) . Other viral proteins influence entry in a context-dependent manner. For example, UL21 is required for the syncytial phenotype associated with gB syncytial mutants but is dispensable for syncytial mutations in gK, UL20, or UL24 (20). In addition, the syncytial phenotypes of gB or gK mutants are reduced in gM- or UL11-negative backgrounds (21).

To examine the HSV-1 entry machinery, we previously generated viruses encoding altered forms of gB. Replacement of HSV-1 gB with SaHV-1 gB in an engineered virus resulted in a small-plaque phenotype (22). Similarly, three alanine substitutions in HSV-1 gB (I671A, H681A, and F683A; gB3A) produced a small-plaque phenotype and delayed penetration into cells (23, 24). Serial passage of both of these fusion-defective viruses selected for second-site mutations that partially restored plaque size (22, 25, 26). These passage experiments identified mutations in the core fusion machinery glycoproteins, demonstrating that *in vitro* selection can reveal alternative routes to efficient entry within the HSV entry machinery.

Notably, serial passage of these viruses also selected mutations in gC (Table 1); however, the role of these gC mutants in entry and fusion was not investigated at the time. These gC mutants are the focus of the present study.

**TABLE 1.**
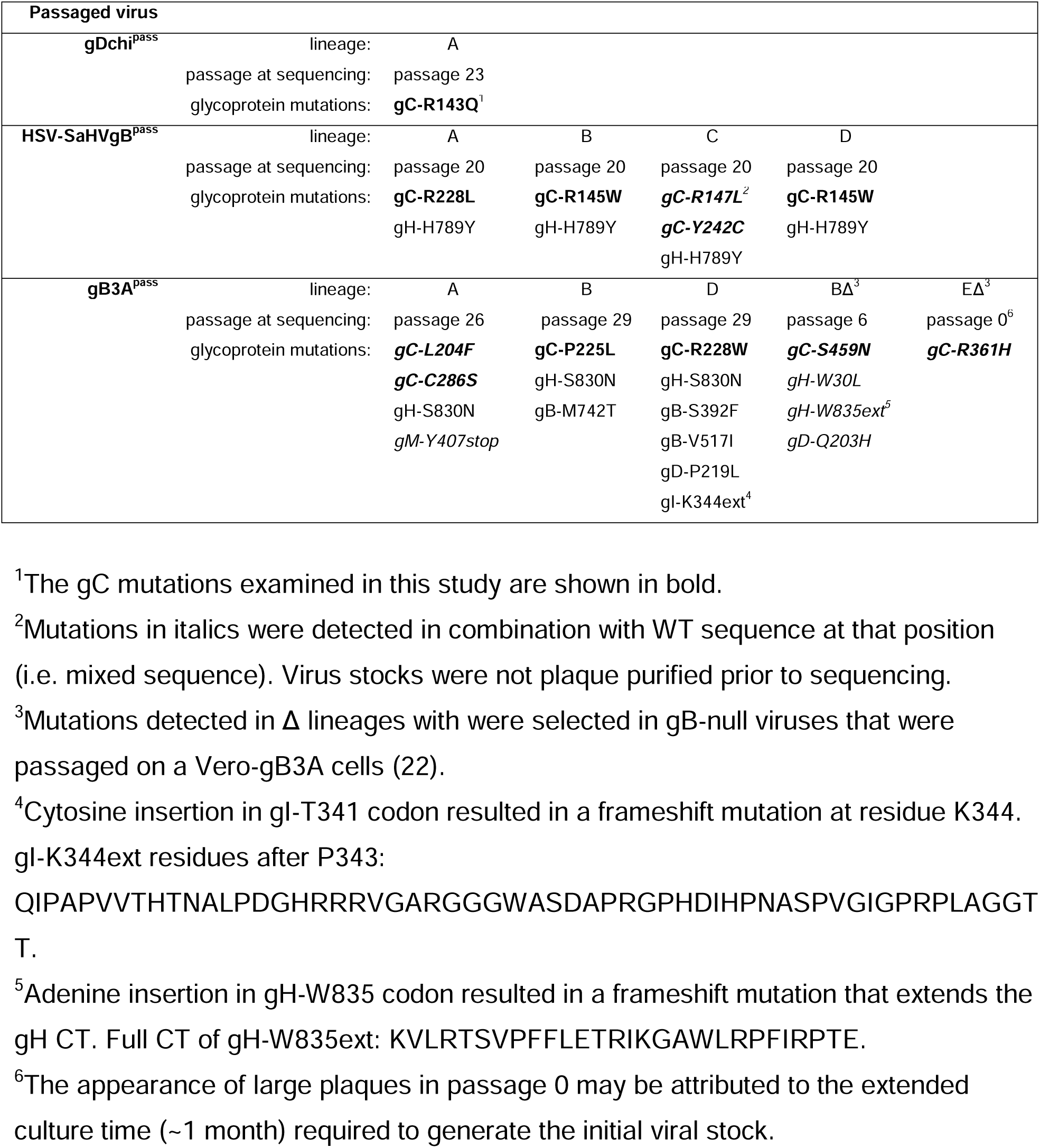
Glycoprotein mutations acquired in passaged virus isolates.

All alphaherpesviruses encode glycoprotein C (gC), an integral membrane glycoprotein that mediates attachment to cell-surface heparan sulfate (27, 28) and contributes to immune evasion by protecting virions from complement-mediated neutralization (29, 30). gC is dispensable for viral entry; however, deletion of gC reduces cell binding and infectivity (31–34). Early studies mapped a macroplaque phenotype to a frameshift mutation in gC (35, 36). Recent studies have shown that deletion of gC reduces entry into cells that support an endocytic route of entry (37–39) and that HSV-1 lacking gC accumulates within endosomes, consistent with a role for gC in penetration from endocytic compartments (40). Intriguingly, the presence of gC modulates the pH sensitivity of gB (37–39) and shields gB from antibody neutralization (37–39), suggesting an interaction between gC and the core fusion machinery.

Together, these findings suggest that gC promotes efficient entry, despite the fact that expression of gC did not enhance fusion mediated by gD, gB, and gH/gL in a cell-cell fusion assay (40).

This study examines the contribution of the selected gC mutations to HSV-1 plaque morphology, entry, and cell-cell fusion using a gD chimeric virus, serial passage, and complementary cell-based assays.

## RESULTS

### Virus carrying HSV-1 gDchi has small plaques

Our previous study demonstrated that cell fusion was impaired when the “profusion domain” (PFD) of HSV-1 gD (residues L268-Y306) was replaced by the corresponding region of SaHV-1 gD (residues E272-P307) (6). The results suggested that this region contributed to a species-specific interaction between the gD and gH. To further investigate the functional interaction of this gD chimera with other glycoproteins, we generated and characterized HSV-1 virus carrying the chimera (pQF137, designated HSV-1 gDchi in this study). We created an HSV-1 BAC carrying gDchi by first deleting US6 (the gene encoding gD) from the HSV-1 BAC pGS3217 and then recombining the HSV-1 gDchi into the BAC. Intermediate BACs (with kan^R^ insertions) and final BACs (with the kan^R^ removed) were confirmed by at least four restriction enzyme digestions.

HSV-1 gDchi or WT (pGS3217) BAC DNA was transfected into Vero cells expressing Cre recombinase to excise the BAC backbone, and virus was harvested from the cells. Vero cells were infected with samples from these transfections to generate WT (GS3217) and HSV-1 gDchi virus stocks. Vero cells were infected with HSV-1 gDchi and WT virus at 0.01 pfu/cell. Due to the growth defects of HSV-1 gDchi virus, infected cells and supernatants were harvested 7-14 days post-infection for the HSV-1 gDchi virus and 3 days post-infection for the WT virus.

Interestingly, like gB3A viruses and HSV-SaHVgB viruses, HSV-1 gDchi virus also displays a small plaque phenotype (Fig.1A). Compared to the WT (GS3217) plaques, the HSV-1 gDchi plaques were approximately 200-fold smaller (Fig. 1A).

**Figure 1.**
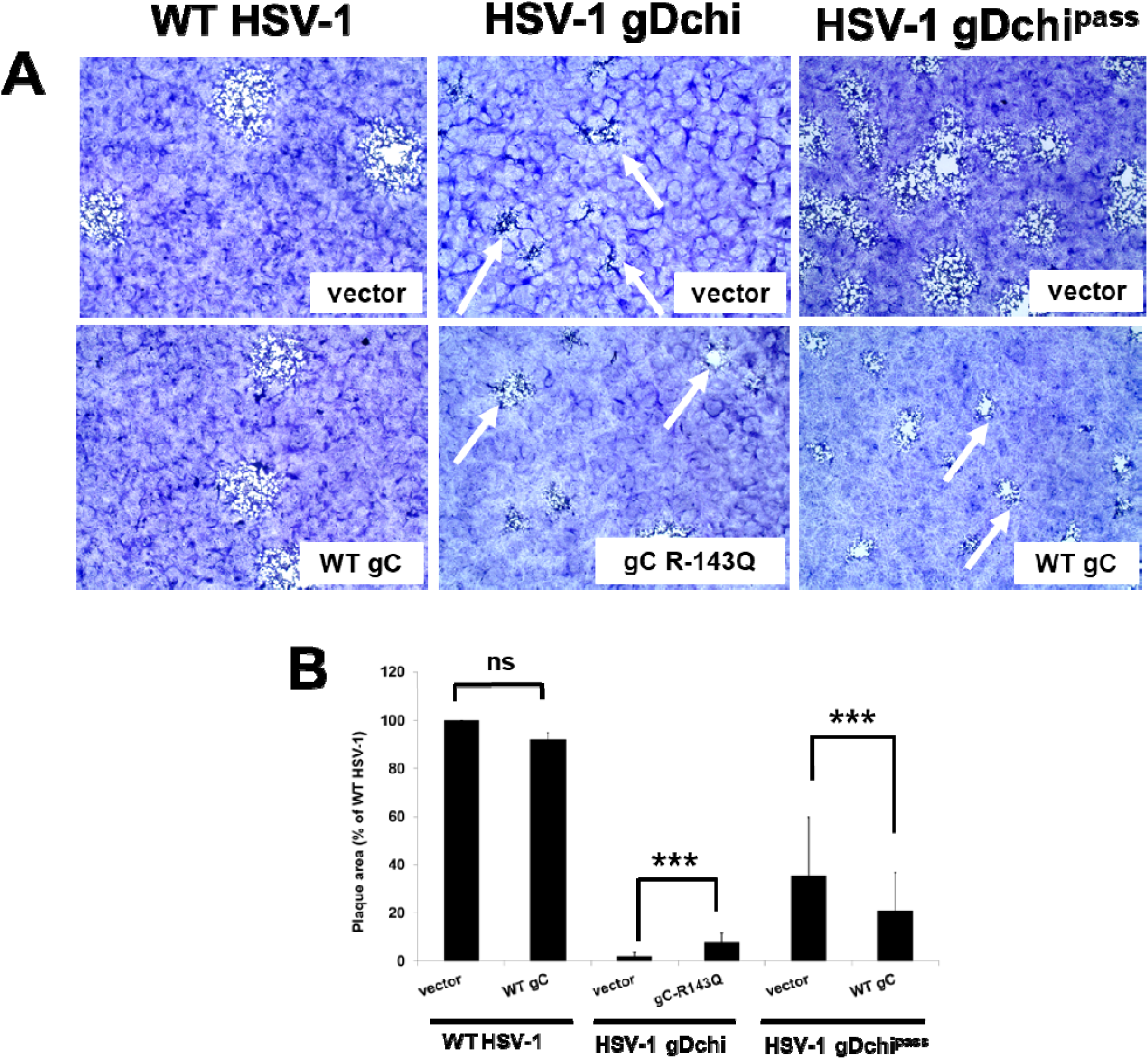
HSV-1 gDchi plaque size is partially restored by a gC mutant. (A) Vero cells were transfected overnight with WT gC, gC-R143Q, or an empty vector. Cells were then infected with WT HSV-1, HSV-1 gDchi, or a passaged isolate of gDchi (gDchi^pass^) at 150 PFU/well. Cells were stained with Giemsa stain three days post-infection and plaques were imaged at 40X magnification. (B) The average plaque area was calculated and expressed as a percentage of the average plaque area for WT HSV-1 on vector-transfected cells. Error bars represent standard deviations. Asterisks indicate a significant difference (Mann-Whitney U test, *** *p* ≤ 0.001), and ns indicates not significant (*p* > 0.05).

### Serially passage of HSV-1 gDchi selects for a mutation in gC

We used serial passage to select for second-site mutations that would rescue the HSV-1 gDchi plaque morphology. We hypothesized that the location of these second-site mutations would reveal sites of functional importance on gD or gH. HSV-1 gDchi virus harvested from Vero-Cre cells was used to select for revertant viruses possessing a growth advantage. The HSV-1 gDchi virus stocks were passaged serially in Vero cells using a multiplicity of infection (MOI) of 0.01. Virus was titered after each passage. After serial passage, larger plaques were observed during titration and the infections spread faster in culture than the parental HSV-1 gDchi virus. By passage 23, plaque areas from the passaged virus stock (HSV-1 gDchi^pass^) were larger than (10-100 times) the HSV-1 gDchi virus (p ≤ 0.01) (Fig. 1B). We sequenced two independent isolates of HSV-1 gDchi^pass^ and unexpectedly found no mutations in HSV-1 gDchi, gB, gH or gL. The only glycoprotein mutation was gC-R143Q, which emerged in two gDchi^pass^ lineages.

### gC-R143Q contributes to the restoration of plaque size after passage of HSV-1 gDchi

To investigate whether gC-R143Q facilitated partial restoration of plaque size, we generated plasmids encoding WT gC (pQF463) and gC-R143Q (pQF465). We transfected Vero cells with gC-R143Q overnight, infected the cell with HSV-1 gDchi, and measured plaque size three days post-infection (Fig. 1). HSV-1 gDchi plaque size was significantly enhanced by expression of gC-R143Q (4.5-fold, p≤0.01) (Fig. 1B).

To further examine whether gC affected HSV-1 gDchi plaque size, we determined whether exogenous expression of WT gC would suppress the enhanced plaque size of HSV-1 gDchi^pass^ by competing with gC-R143Q. We transfected Vero cells with WT gC overnight, infected the cells with HSV-1 gDchi^pass^ or WT HSV-1, and measured plaque sizes three days post-infection (Fig. 1). For WT virus infections, transfection with gC did not change plaque size (Fig. 1B). In contrast, HSV-1 gDchi^pass^ plaque size was significantly smaller on gC-transfected cells compared to vector-transfected cells (p ≤ 0.01) (Fig. 1B), suggesting that gC-R143Q contributes to the increased plaque size observed in HSV-1 gDchi^pass^. Given these results and the fact that HSV-1 gDchi^pass^ does not carry mutations in the core fusion machinery, we concluded that gC-R143Q likely contributes to increased plaque size of HSV-1 gDchi^pass^ viruses.

### gC mutations acquired during passage of mutant gB viruses contribute to the restoration of plaque size observed after passaging

We previously demonstrated that HSV-SaHVgB and gB3A viruses have small plaque phenotype and serial passage of these viruses selected for revertant viruses (HSV-SaHVgB^pass^ and gB3A^pass^) with larger plaque sizes (**22, 25, 26**). We determined that mutations acquired in gH were partially responsible for the increased plaque size of these passaged viruses (**22**). Interestingly, HSV-SaHVgB^pass^ virus isolates acquired several mutations in gC in addition to the mutation gH-H789Y (Table 1). Similarly, the gB3A^pass^ virus isolates acquired several gC mutations in addition to mutations in gH (gH-S830N or W835ext).

We investigated whether these gC mutations contributed to the restoration of plaque size for HSV-SaHVgB^pass^ and gB3A^pass^ using the same approach we applied to gDchi^pass^. To test whether exogenous expression of WT gC would reduce plaque size of the passaged viruses by competing with the gC mutants, we transfected the Vero cells with WT gC overnight, infected the cells with WT, HSV-SaHVgB^pass^ or gB3A^pass^ viruses, and measured plaque sizes. For WT virus, plaque size was unchanged by expression of gC (Fig. 2). In contrast, plaque sizes of HSV-SaHVgB^pass^ or gB3A^pass^ viruses were significantly reduced in gC-transfected cells compared to vector-transfected cells. These results indicate that WT gC expression suppressed the plaque size of the revertant viruses, presumably by competing with the gC mutants present in these passaged viruses.

**Figure 2.**
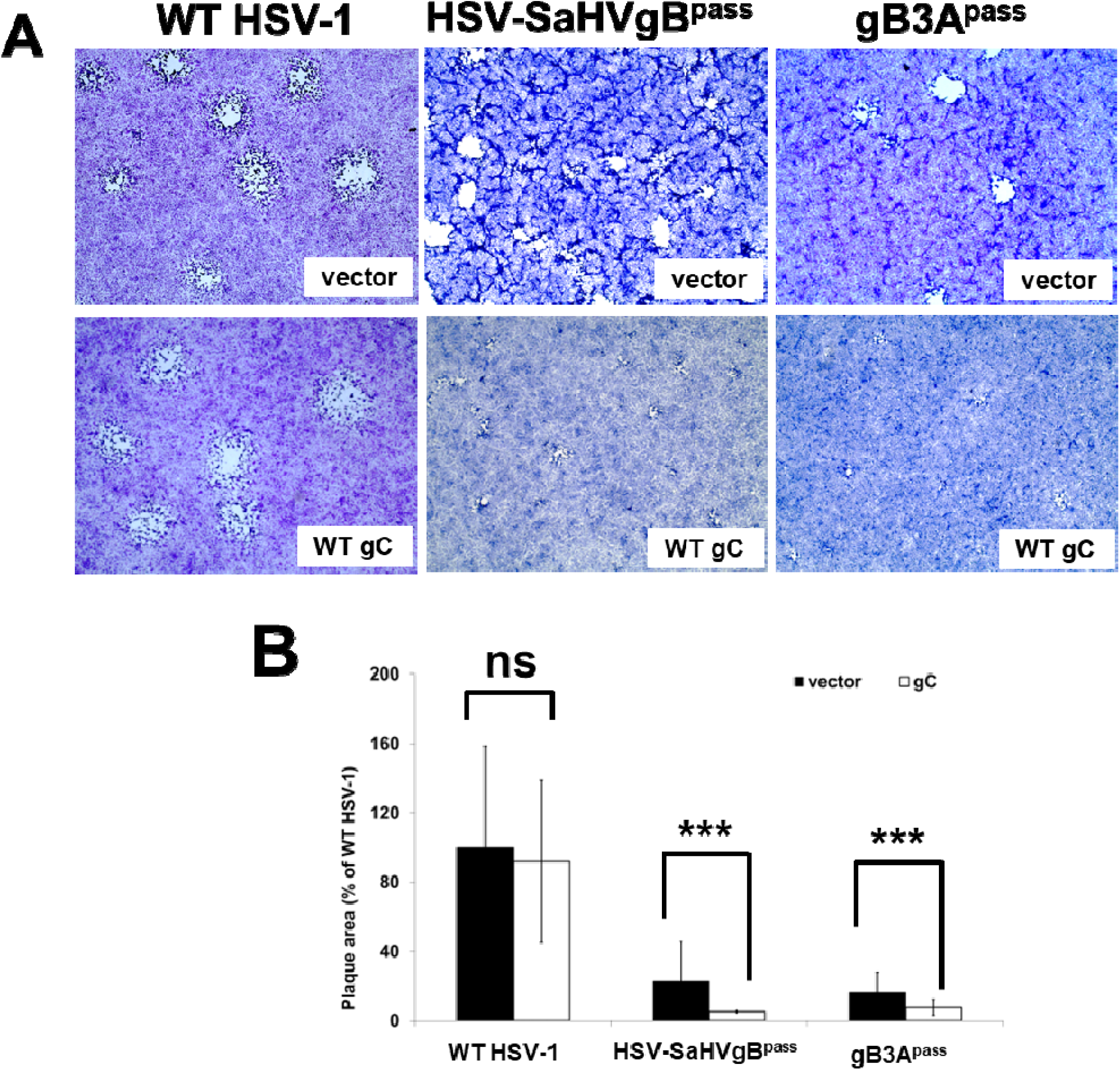
Overexpression of WT gC reduces plaque size of passaged gB mutant viruses. (A) Vero cells were transfected overnight with WT gC or an empty vector. Cells were then infected with a passaged isolate gB3A (gB3A^pass^) or a passaged isolate of HSV-SaHVgB (HSV-SaHVgB^pass^) at 150 PFU/well. Three days after infection, cells were stained with Giemsa and plaques were imaged at 40X magnification. (B) The average plaque area was calculated and expressed as a percentage of the average plaque area for WT HSV-1 on vector-transfected cells. Error bars represent standard deviations. Asterisks indicate a significant difference (Mann-Whitney U test, *** *p* ≤ 0.001), and ns indicates not significant (*p* > 0.05).

To further examine the contribution of gC to plaque size, we determined the effect of exogenous expression of gC mutants on the plaque size of the mutant gB viruses prior to passage. In our previous study, gC-L204F and gC-R228W were selected after passage of gB3A virus, whereas gC-R145W and gC-R228L were selected after passage of HSV-SaHVgB virus (**22**). Thus, Vero cells were transfected with gC-L204F, gC-R228W prior to infection with the gB3A virus (Fig. 3A), or with gC-R145W orgC-R228L prior to infection with the HSV-SaHVgB virus (Fig. 3B). Plaques were measured three days post-infection. Expression of gC-L204F or gC-R228W significantly increased gB3A virus plaque size (p≤0.01, Fig. 3C), and expression of gC-R145W or gC-R228L significantly increased HSV-SaHVgB virus plaque size (p≤0.01, Fig. 3D). Overall, expression of these gC mutants increased plaque size for the gB mutant viruses by approximately 1.4- to 2.5-fold.

**Figure 3.**
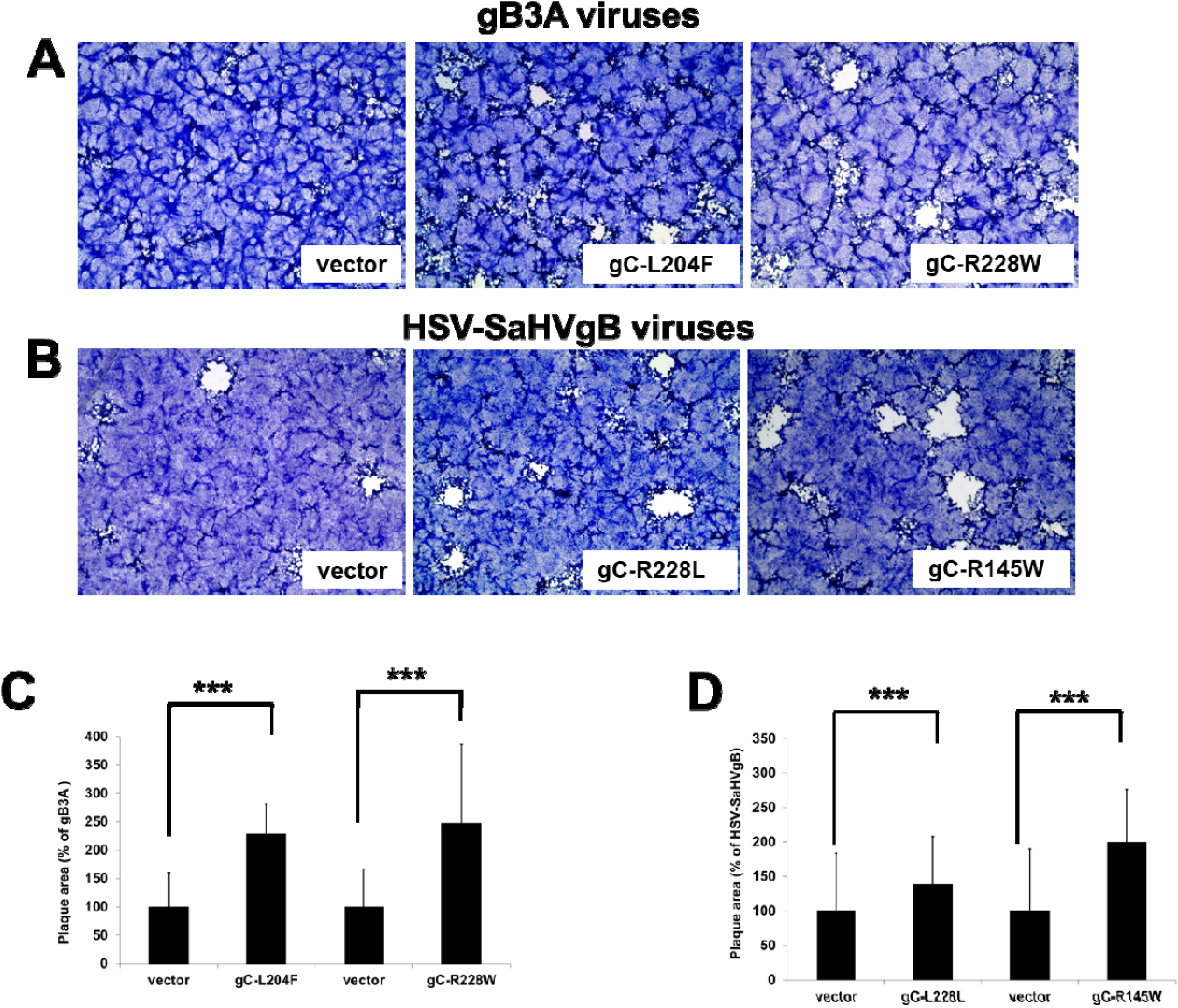
HSV-1 gB3A and HSV-SaHVgB plaque size is partially restored by gC mutants. (A) Vero cells were transfected overnight with a gC mutant or an empty vector. Cells were then infected with gB3A virus (A) or HSV-SaHVgB virus (B) at 150 PFU/well. After three days, cells were stained with Giemsa and plaques imaged at 40X magnification. Average plaque areas for gB3A virus (C) or HSV-SaHVgB virus (D) were calculated and expressed as a percentage relative to plaques generated by those viruses on vector-transfected cells. Error bars represent standard deviations. Asterisks indicate a significant difference (Mann-Whitney U test, *** *p* ≤ 0.001), and ns indicates not significant (*p* > 0.05).

Taken together, the enhancement of plaque size for these gB mutant viruses upon exogenous expression of mutant gC (Fig. 2), and the suppression of plaque size for the passaged gB mutant viruses upon expression of WT gC (Fig. 3), suggest that gC contributes to the regulation of plaque size.

### gC mutants do not enhance WT cell-cell fusion

Although HSV-1 entry and fusion require four essential glycoproteins (gD, gB, and gH/gL), and gC is not critical for this process, the observed effects of gC on plaque size in these mutant viruses suggest that gC modulates fusion activity. To directly assess the effect of gC on fusion, we employed a cell-cell fusion assay. We generated expression constructs to test eleven gC mutants selected following passage of gDchi, gB3A, or HSV-SaHVgB viruses (Table 1). Cell surface expression of the gC mutants was assessed by cell-based ELISA (CELISA) in transfected CHO-K1 cells. WT gC was readily detected on the cell surface (Fig. 4A). Most gC mutants (gC-R143Q, gC-R145W, gC-R147L, gC-P225L, gC-R228L, gC-C286S, gC-R361H and gC-S459N) exhibited expression levels comparable to WT gC, suggesting that these proteins were properly folded and transported to the plasma membrane. In contrast, some gC mutants showed reduced surface expression, including gC-R228W, gC-Y242C, and gC-L204F (p≤0.05), indicating that these mutants may not be properly processed.

**Figure 4.**
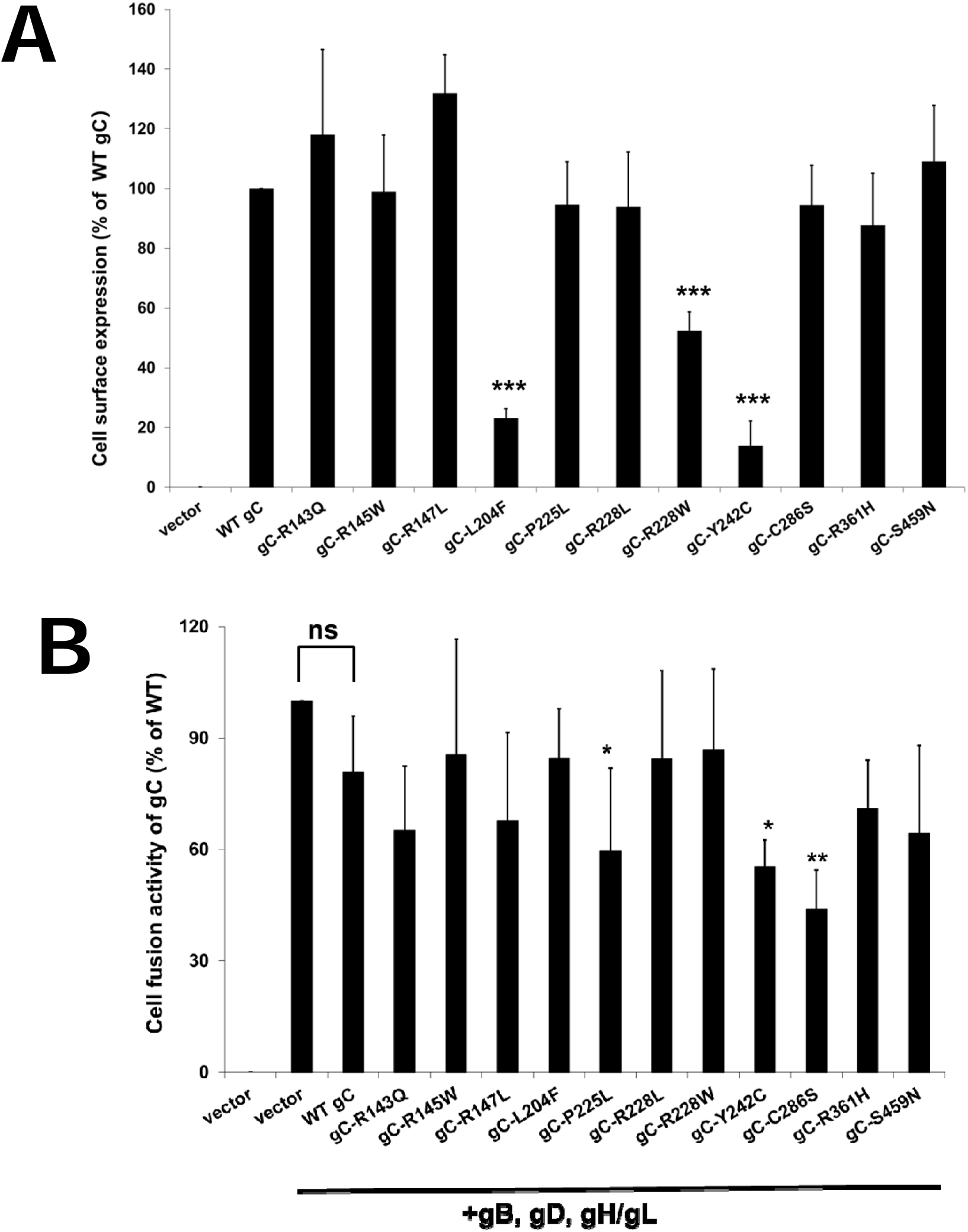
gC mutants do not enhance WT cell-cell fusion. (A) Cell surface expression of gC mutants. CHO-K1 cells were transfected in 96-well plates with WT gC, gC mutants, or empty vector (pSG5). After rinsing, gC expression was detected by CELISA using an anti-HSV-1 gC monoclonal antibody. Error bars represent standard deviations. (B) Cell-cell fusion activity. CHO-K1 cells (targets) were transfected with nectin-1 and a reporter plasmid encoding luciferase under the control of the T7 promoter. A second set of CHO-K1 cells (effectors) were transfected with T7 polymerase, gB, gD, gH, and gL, plus vector, WT gC or mutant gC. After 18 h, effector target and effector cells were mixed. After six hours, cells were lysed and luciferase activity was quantified as a measure of cell fusion. Each bar shows the means of at least three independent determinations. The results are expressed as a percentage of WT HSV-1 activity in the absence of gC after subtraction of the background values. The error bars represent standard deviations. Asterisks indicate a significant difference (independent Student’s t-test, *** *p* ≤ 0.001, ** *p* ≤ 0.01, * *p* ≤ 0.05) and ns indicates not significant.

To assess the function of these gC mutants in cell-cell fusion, a set of CHO-K1 cells (effector cells) was transfected with T7 polymerase, gD, gB, gH/gL and either WT gC or a gC mutant. These effector cells were cocultured with a set of CHO-K1 cells transfected with nectin-1 receptor and a plasmid encoding luciferase under the control of a T7 promoter. Luciferase activity was used as a quantitative measure of cell-cell fusion.

WT gC co-transfected with gB, gD, and gH/gL did not significantly affect fusion activity compared to cells transfected with gB, gD, and gH/gL and vector (Fig. 4B), consistent with a recent study (40). Similarly, none of the gC mutants enhanced fusion. The results indicate that none of the gC mutations selected by passage of our mutant viruses are globally hyperfusogenic. In fact, some of the gC mutants inhibit fusion, which could result from non-functional interactions with WT gD, gH/gL, and/or gB that interfere with fusion. Alternatively, expression of these gC mutants may impact expression of the WT entry glycoproteins, resulting in fusion.

### gC mutants do not enhance WT HSV-1 entry

Although the cell-cell fusion assay isolates the functions of the core entry glycoproteins during viral entry, HSV-1 virions contain additional glycoproteins that may influence entry and spread. Therefore, we used a complementation assay to determine whether gC mutants affect virus entry when all viral glycoproteins are present. Vero cells were transfected with WT gC or a gC mutant and subsequently infected with gC-null HSV-1 (41). After an overnight incubation, virus particles carrying exogenously expressed WT or mutant gC were harvested. To determine the impact of the gC mutants on virus entry, the complemented viruses were titered on Vero cells expressing WT gC. Higher titers indicate increased levels of viral entry. The titers of virus carrying WT gC or gC mutants were not significantly different (Fig. 5A). Together, these results suggest that gC mutations do not significantly impact virus entry in the presence of WT glycoproteins.

**Figure 5.**
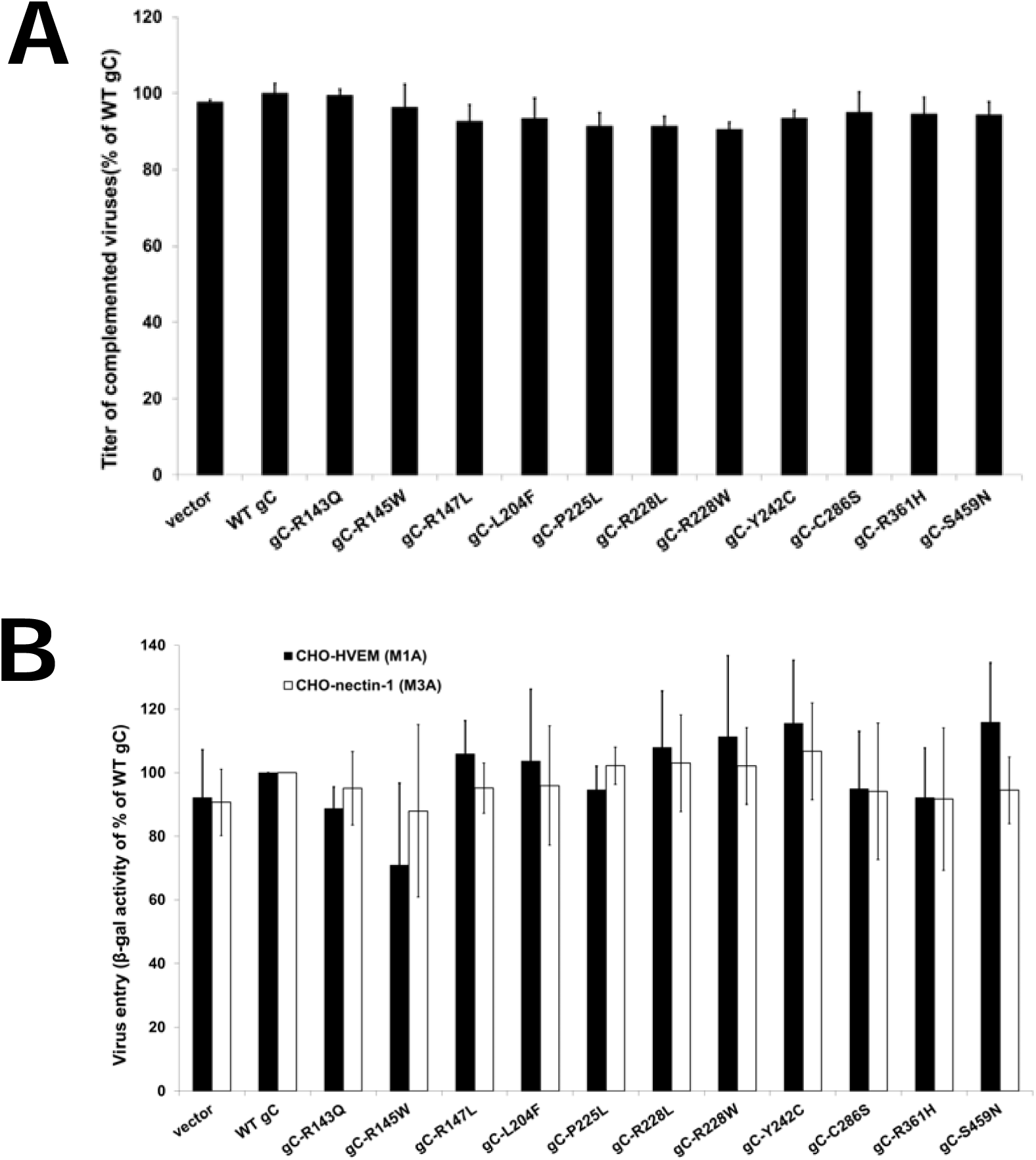
gC mutants do not enhance virus entry. (A) Complementation of gC-null virus by gC mutants. Vero cells were transfected with plasmids encoding WT gC, mutant gC, or empty vector. After overnight incubation, cells were infected with gC-null HSV-1 (41). Virus stocks, consisting of virions phenotypically complemented with mutant forms of gC, were harvested 24 h post-infection and titered on Vero cells to assess relative efficiency of entry. Results are representative of three independent experiments. (B) CHO-K1 cells that carry *lacZ* under control of the ICP4 promoter and express HVEM (M1A) or nectin-1 (M3A) were incubated with dilutions gC-null virus that was complemented with WT gC, gC mutants, or empty vector (pSG5). After five hours at 37°C, β-galactosidase activity was quantified as a measure of virus entry. The data are presented as percentage signal detected for gC-null virus complemented with WT gC. Results are representative of three independent experiments.

To confirm these findings, we performed an entry assay using CHO cells that express one of two gD receptors, nectin-1 (M3A cells) or HVEM (M1A cells) and carry the *lacZ* reporter gene under the control of the HSV ICP4 promoter. Cells were infected with gC-null HSV-1 viruses that had been complemented with the gC mutant, and viral entry was quantified by measuring β-galactosidase activity, as previously described (42, 43). No significant differences in viral entry into HVEM- or nectin-1-expressing cells were observed among cells infected with virus complemented with WT gC or any of the gC mutants (Fig. 5B). These results confirm that these gC mutants do not impact viral entry in the presence of WT glycoproteins.

### WT gC and gC mutants enhance cell-cell fusion when coexpressed with a corresponding gH mutant

Although the gC mutants did not enhance cell-cell fusion when WT gD, gH/gL, and gB were expressed, given that gC mutants enhanced plaque size of mutant viruses (Fig. 1 and 3), we investigated whether the gC mutants would enhance fusion in the cell-cell fusion assay when mutant viral glycoproteins were present. We limited the gC mutants examined to gC-R145W and gC-R228L, which emerged with gH-H789Y in HSV-SaHVgB^pass^ viruses, as well as gC-P225L and gC R228W, which emerged with gH-S830N in gB3A^pass^ viruses (**22**).

We first tested whether WT gC or gC mutants affected cell-cell fusion mediated by gH-H789Y and gH-S830N mutants when co-expressed with WT gL, gD, and gB (Fig. 6A). As shown previously, both gH-H789Y and gH-S830S display reduced fusion compared to WT gH (22). Remarkably, WT gC enhanced fusion significantly when gH-H789Y or gH-S830S was expressed, in contrast to the previous results with all WT entry glycoproteins (Fig. 4). These results indicate that WT gC partially compensates for the fusion defects of the gH mutants.

**Figure 6.**
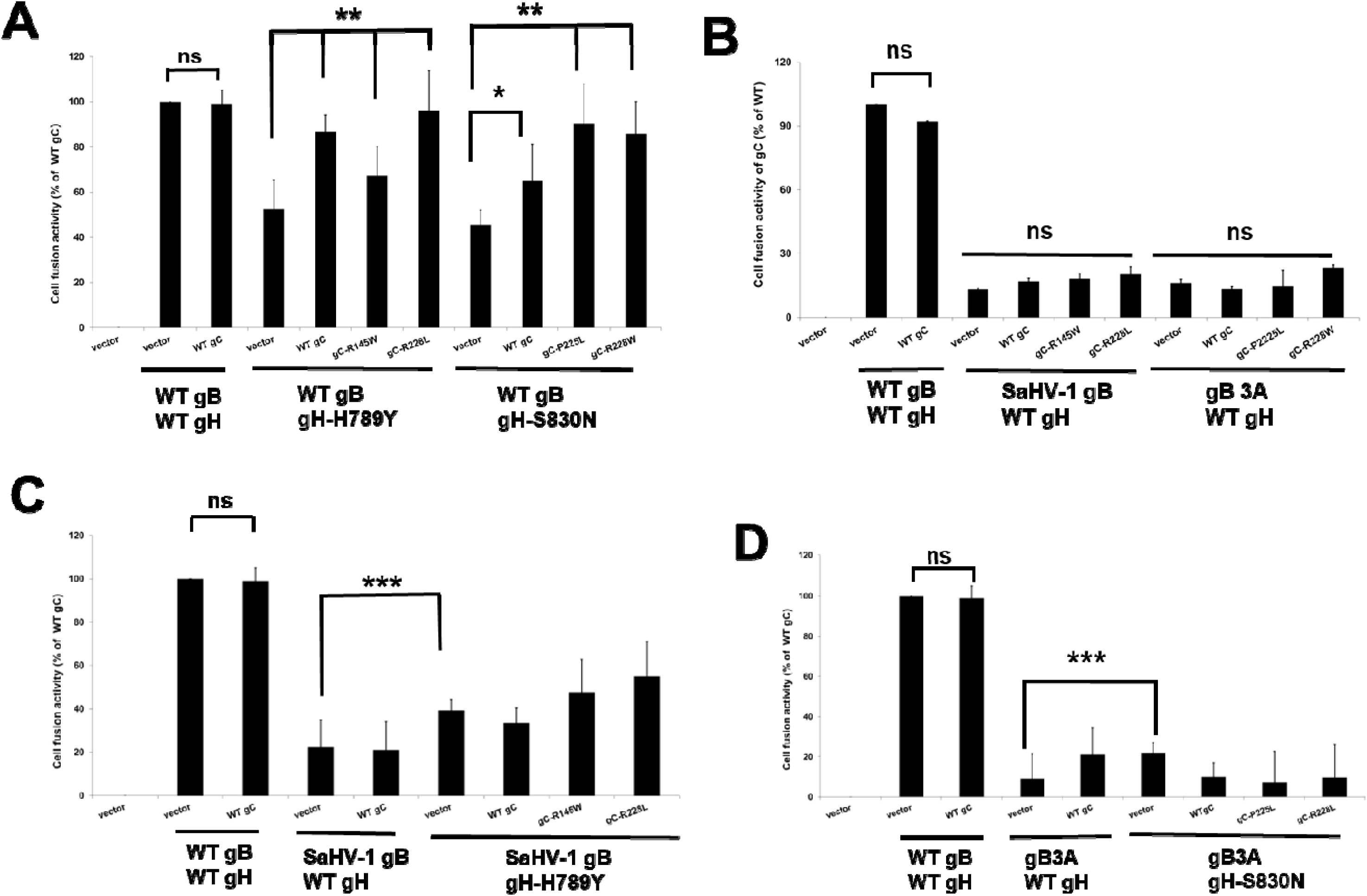
gC enhances cell-cell fusion mediated by deficit gH mutants. CHO-K1 cells (targets) were transfected with nectin-1 and a reporter plasmid encoding luciferase under the control of the T7 promoter. A second set of CHO-K1 cells (effectors) were transfected with T7 polymerase, gD, gL, WT or mutant gB, WT or mutant gH, plus vector, WT gC or mutant gC. After 18 h, effector target and effector cells were mixed. After six hours, cells were lysed and luciferase activity was quantified as a measure of cell fusion. Each bar shows the means of at least three independent determinations. The results are expressed as a percentage of WT HSV-1 activity in the absence of gC after subtraction of the background values. The error bars represent standard deviations. Asterisks indicate a significant difference (independent Student’s t-test, *** *p* ≤ 0.001, ** *p* ≤ 0.01, * *p* ≤ 0.05) and ns indicates not significant. (A) Impact of gC mutants on fusion mediated by gH mutants. (B) Impact of gC mutants on fusion mediated by gB mutants. (C) Impact of gC mutants on fusion mediated by SaHV-1 gB and gH-H789Y. (D) Impact of gC mutants on fusion mediated by gB3A and gH-S830N.

In addition to WT gC, the gC mutants also significantly increased fusion mediated by gH-H789Y or gH-S830N (p ≤ 0.01). In fact, three of the four gC mutants enhanced fusion to a greater extent than did WT gC when coexpressed with gH-H789Y or gH-S803N, resulting in near-WT fusion levels. These results suggest that the gC can enhance cell-cell fusion, but this effect is only detected when cell-cell fusion is operating at sub-optimal levels due to a deficient gH.

### WT gC and gC mutants fail to enhance cell-cell fusion when coexpressed with a corresponding gB mutant

We next tested whether WT gC or gC mutants enhance fusion mediated by gB3A or SaHV-1 gB. The low levels of fusion mediated by these forms of gB were not rescued by the expression of WT gC or gC mutants (Fig. 6B).

We then evaluated whether WT gC and gC mutants affect fusion activity when coexpressed with both their corresponding gH mutants and the corresponding SaHV-1 gB or gB3A mutant (i.e. the version of gB present in their parental viruses). As shown previously, gH-H789Y significantly enhanced fusion when coexpressed with SaHV-1 gB relative to WT gH (Fig. 6C). Similarly, gH-S830N enhanced fusion when coexpressed with gB3A relative to WT gH (Fig. 6D) (22). Unexpectedly, neither WT gC nor gC mutants further increased fusion activity when coexpressed with matched gB-gH mutant pairs (Fig. 6C and 6D). These results suggest that WT gC and gC mutants cannot rescue the impaired fusion observed when both gB and gH are mutated. Despite the fact that the gC mutants impact plaque size (Fig. 1 and 3) and enhance fusion mediated by the gH mutants (Fig. 6A), a direct cooperation among gC, gH, and gB mutants was not detected by the cell-cell fusion assay.

## DISCUSSION

### Established roles for gC in HSV-1 entry and pathogenesis

In addition to the core fusion machinery of gD, gH/gL, and gB, HSV-1 encodes more than a dozen envelope glycoproteins, several of which can influence entry and fusion under specific conditions. In addition to mediating immune evasion through complement binding (44–51), gC has been shown to modulate entry and fusion. HSV-1 gC binds to heparan sulfate on the cell surface (45–52). Although gC is dispensable for entry, deletion of gC reduces cell binding and infectivity (32, 53), and anti-gC antibodies inhibit both cell binding and HSV infection (32, 53). In fact, gC is a target of antibody neutralization (54, 55) and may partially shield the core fusion machinery from antibody neutralization (37–39, 56), suggesting that gC is in close proximity to gD, gH/gL, and/or gB.

Accumulating evidence suggests that gC promotes HSV-1 entry at a step beyond initial cell attachment. In cell types that support an endocytic route of entry, deletion of gC reduces viral entry and the presence of gC modulates the sensitivity of gB to acidic pH, increasing the pH threshold required to detect antigenic changes in gB (37–39). HSV-1 lacking gC is endocytosed at normal levels but accumulates in the endosome, suggesting that gC contributes to penetration from the endosome during entry (40). Moreover, incorporation of gC into a pseudotyped virus carrying the HSV-1 core fusion machinery enhanced viral entry in a cell-type-specific manner, including in HaCaT cells, which support an endocytic route of entry (57). These findings suggest that gC may functionally interact with the core fusion machinery.

*In vivo*, the contribution of gC to HSV-1 virulence depends on the experimental system. Although deletion of gC had no impact on HSV-1 virulence in mice (58) and an HSV-2 virus encoding a gC mutant retained wild-type virulence after intravaginal or intracerebral inoculation (59), gC contributed to virulence in other models. Glycoprotein C supported efficient replication in human skin in a SCID-hu mouse model (49, 60–62), enhanced viral replication in mouse models with intact complement-mediated immunity (49, 60–62), and increased replication and neurovirulence following intranasal infection in a rabbit model (49, 60–62).

### Selection of gC mutations in fusion-defective viruses

Building on these observations, our study further supports a role for gC as a regulator of HSV-1 entry that interacts functionally with the core fusion machinery. Several of the gC mutants in this study were identified previously after serial passage of two HSV-1 engineered to express mutant gB (gB3A and HSV-SaHVgB). Both viruses exhibited a small-plaque phenotype that was partially rescued after serial passage (22, 24). Passaged virus isolates acquired multiple mutations in gH and gC (Table 1), consistent with prior work showing that serial passage can select extragenic suppressor mutations in HSV(63). We previously demonstrated that the gH mutations contributed to the partial restoration of plaque size, but the role of the gC mutations in restoration of plaque size was not investigated.

Leveraging natural selection in a similar manner, we generated a mutant HSV-1 encoding a chimeric version of gD (gDchi) for this study. Similar to the mutant gB viruses, gDchi exhibited a small plaque phenotype that was partially rescued after passage (Fig. 1), and whole genome sequencing revealed a mutation in gC (Table 1).

### gC as a conditional regulator of HSV-1 entry

To examine whether these gC mutations contributed to plaque size restoration, we infected Vero cells expressing WT gC with the passaged viruses, allowing WT gC to compete with mutant gC. Expression of WT gC reduced plaque size of the passaged viruses (Fig. 1 and 2), indicating that gC mutations contributed to the enlarged plaque phenotype. Moreover, expression of mutant gC enhanced plaque size of viruses prior to passage (Fig. 1 and 3), providing more direct evidence that gC mutations promote increased plaque size.

To assess their effects on entry and fusion, we generated expression constructs for a panel of gC mutants (Fig. 4A). Although these gC mutations did not enhance fusion mediated by the wild-type core fusion machinery (Fig. 4B) or the entry of gC-null virus (Fig. 5), they were repeatedly selected during serial passage of fusion-defective viruses (Table 1). These findings indicate that HSV-1 gC functions as a conditional accessory regulator that can partially compensate for defects in the core fusion machinery. Enhancement of cell-cell fusion by gC was observed only when the entry machinery was operating at a suboptimal level due to mutations in gH (Fig. 6A). This context-dependent contribution to fusion may explain why gC mutations arose during passage of all three distinct fusion-defective viruses, including HSV-SaHVgB, gB3A, and gDchi.

HSV-1 initially attaches to the cell surface via gC- and gB-mediated heparan sulfate interactions. Following attachment, gD binds receptor and undergoes conformational changes that signal gH/gL. Structural and biochemical studies have demonstrated receptor-induced conformational changes in gD(1) and direct binding of gD and gH/gL (64, 65). We previously identified a species-specific functional interaction between gD and gH/gL (6), and the impaired spread of the gDchi virus is likely due to defects in the gD-gH/gL signaling. For gDchi virus, the selection of second-site mutations in gC, rather than in gD or gH/gL, suggests that gC can partially compensate for inefficient gD-gH/gL signaling.

This compensatory role for gC is supported by our analysis of fusion-defective viruses carrying gB mutants. Serial passage of these small-plaque viruses (HSV-SaHVgB or gB3A) selected for second-site mutations in gH (gH-H789Y and gH-S830N, respectively) that partially rescued plaque size (22). Despite the ability of these gH mutants to compensate for fusion defects when expressed with their respective gB mutants, they exhibited reduced fusion activity when co-expressed with wild-type gB (Fig. 6A)(22), demonstrating that the gH mutations introduce functional deficits of their own. The concurrent selection of gC mutations along with these gH mutations during passage suggests that gC may function as a secondary compensatory factor, partially restoring fusion efficiency compromised by the gH mutations. These gC mutants may enhance gD-gH/gL functional interactions, as proposed above for the gDchi virus, since both WT gC and gC mutants increased cell-cell fusion when coexpressed with gH mutants (Fig. 6A). Alternatively, gC may support gB function of gB-gH/gL interaction, since expression of gC mutants enhanced plaque size of the gB-mutant viruses prior to passage, while WT gH was present (Fig. 3).

Additionally, gC mutations may promote fusion by enhancing heparan sulfate binding; however, enhanced fusion was not observed with in a wild-type background (Fig. 4B) suggesting that altered attachment alone would not account for the phenotype. Instead, the functional impact of gC appears to depend on the state of the core fusion machinery, particularly gH and gB. Together, our findings support a model in which HSV-1 gC functions as a conditional accessory regulator that modulates entry and fusion through functional interactions with the core fusion machinery, compensating for defects in gD, gH/gL, or gB signaling under suboptimal conditions.

## MATERIALS AND METHODS

### Cells and antibodies

Chinese hamster ovary (CHO-K1; American Type Culture Collection (ATCC, USA)) cells were grown in Ham’s F12 medium supplemented with 10% fetal bovine serum (FBS) (ThermoFisher Scientific, USA). Vero cells (ATCC, USA) were grown in Dulbecco modified Eagle medium (DMEM) supplemented with 10% FBS, penicillin and streptomycin. M1A and M3A cells (66) were derived from CHO-IEβ8 cells to stably express human HVEM and nectin-1, respectively. These cells were grown in Ham’s F-12 medium supplemented with 10% FCS, 150 μg of puromycin/ml, and 250 μg of G418/mL. Vero-gB3A cells (22) were derived from Vero cells to stably express gB3A and grown in DMEM supplemented with 10% FCS and 250 μg of G418/mL. gC was detected using anti-HSV-1 gC MAb (EastCoast Bio, USA).

### Construction of gDchi BAC

BACs were generated by manipulation of the BAC GS3217, which was kindly provided by Dr. Gregory Smith (Northwestern University) and Dr. Yasushi Kawaguchi (University of Tokyo) (67, 68). GS3217 is an HSV-1 F strain BAC that carries the red fluorescence protein (RFP) tdTomato reporter gene with a nuclear localization signal under the control of a CMV IE promoter. The CMV>NLS-tdTomato>pA cassette is inserted after the start codon of US5 (gJ) and introduces an in-frame stop codon.

To generate the gDchi BAC (pQF415), a gD-null BAC pQF284 was first constructed by deleting US6 (the gD gene) from the BAC GS3217. A kan^R^ gene was PCR-amplified from pGS1439 (kindly provided by Dr. Gregory Smith) using primers that consist of kan^R^ homology (underlined) and US6 (gD) flanking sequence: CCCGATCATCAGTTATCCTTAAGGTCTCTTTTGTGTGGTGCGTTCCGGTATACCCC CCCTTAAT<u>AGGATGACGACGATAAGTAGGG</u> and ATCCCAACCCCGCAGACCTGACCCCCCCGCACCCATTAAGGGGGGGTATACCGG AACGCACCAC<u>CAACCAATTAACCAATTCTGAT</u>. Using a two-step red-mediated recombination strategy (69, 70), this PCR product was recombined into pGS3217 in place of US6 to delete gD. The kan^R^ cassette then was recombined out to generate pQF284.

To introduce the HSV-1/SaHV-1 gD chimera (gDchi) into this gD-null BAC, the N-terminal FLAG tag present in the gDchi expression construct pQF137 (6) was replaced first with the native gD signal sequence, generating plasmid pQF355. This construct encodes HSV-1 gD residues M1-L268 and Y306-Y369, with SaHV-1 gD residues E272-P307 replacing the HSV-1 profusion domain.

The kan^R^ gene was PCR-amplified from pGS1439 using the primers that consist of kan^R^ homology (underlined) and US6 (gD) sequence including a unique SalI restriction site (bold): CCGCAC**GTCGAC**GACGACGTGCCTAACGGCGTCGGCCCCACGCGGATCCCAGAG CGGATCGGGC<u>AGGATGACGACGATAAGTAGGGAT</u> and GCACGTCGTC**GTCGAC**<u>CAACCAATTAACCAATTCTGATTAGA</u>. This PCR product was digested with SalI and ligated into SalI digested pQF355, The FLAG-tag signal sequence in pQF137 was replaced with the native gD signal sequence to generate pQF413.

The gDchi gene containing a kanR insert was PCR-amplified from pQF413 using primers CGAACGACCAACTACCCCGATCATCAGTTATCCTTAAGGTCTCTTTTGTGTGGTGCGTTCCGGTATGGGGGGGGCTGCCGCCAGGTTG and TGGAGTTAAGGTCCCATCCCAACCCCGCAGACCTGACCCCCCCGCACCCATTAAGGGGGGGTATCTAGTAAAACAAGGGCTGGTGCGA. Using two-step red-mediated recombination, this PCR product was recombined into the gD-null BAC pQF284 and then the kanR cassette was recombined out to generate the BAC pQF415, which encodes gDchi in place of WT gD.

At each step of the BAC constructions, the intermediate BACs with kanR insertions and the final BACs were confirmed by at least four restriction enzyme digestions. Sequencing was conducted by the Northwestern Genomic core facility.

### Viruses

WT virus was generated using GS3217 BAC. gB3A virus (24), HSV-SaHVgB virus, gB3A^pass^ viruses, HSV-SaHVgB^pass^ viruses (22), and gC-null virus (HSV-1 ΔgC2-3, KOS strain) (41) have been described previously. To generate a gDchi virus stock, BAC pQF415 was transfected into Vero cells expressing Cre recombinase using Lipofectamine 2000 (Invitrogen, Carlsbad, CA). The transfected cells were harvested three weeks after transfection and sonicated, and the released virus was subsequently passaged on Vero cells to generate gDchi virus stocks.

### Plasmids

Plasmids encoding HSV-1 (KOS strain) gB (pQF439), gD (pQF440), gH (pQF441) and gL (pQF442) (22), as well as human nectin-1 (pBG38) (71), gB3A (23), SaHV-1 gB (6), gH-S830N (pQF444), and gH-H789Y (pQF445) (22) were described previously. pT7EMCLuc plasmid encoding a firefly luciferase reporter gene under the control of the T7 promoter and pCAGT7 plasmid encoding T7 RNA polymerase (72, 73) were used in the fusion assay.

A WT gC (KOS strain) expression construct (pQF463) was generated by subcloning gC from pPEP21 (kindly provided by Dr. Patricia Spear at Northwestern University) into pSG5 vector. Plasmids encoding the gC mutant panel (pQF465-pQF475) were then generated by site-specific mutagenesis of pQF463. The gC mutant plasmids were generated using the following primer pairs: pQF465 (gC-R143Q) with primers GGGTGCAGATCCAATGCCGGTTTCG and CGAAACCGGCATTGGATCTGCACCC. pQF466 (gC-R145W) with primers GCAGATCCGATGCTGGTTTCGGAATTCC and GGAATTCCGAAACCAGCATCGGATCTGC. pQF467 (gC-R147L) with primers GATGCCGGTTTCTGAATTCCACCCGC and GCGGGTGGAATTCAGAAACCGGCATC, pQF468 (gC-L204F) with primers ACCCCCACGTGTTCTGGGCGGAGGG and CCCTCCGCCCAGAACACGTGGGGGT, pQF469 (gC-P225L) with primers ACCGGGCCGCTGCTGACCCAGCGGC and GCCGCTGGGTCAGCAGCGGCCCGGT, pQF470 (gC-R228L) with primers CTGCCGACCCAGCTGCTGATTATCG and CGATAATCAGCAGCTGGGTCGGCAG, pQF471 (gC-R228W) with primers CTGCCGACCCAGTGGCTGATTATCG and CGATAATCAGCCACTGGGTCGGCAG, pQF472 (gC-Y242C) with primers CCCAGGGAATGTGTTACTTGGCCTGG and CCAGGCCAAGTAACACATTCCCTGGG, pQF473 (gC-C286S) with primers CAAGGCGACGTCCACGGCCGCCGCC and GGCGGCGGCCGTGGACGTCGCCTTG, pQF474 (gC-R361H) with primers ACGTTCTCGCGACACAATGCCACCG and CGGTGGCATTGTGTCGCGAGAACGT, pQF475 (gC-S459N) with primers TTCTAGAGCACCACGGCAATCACCAGCCCC and GGGCTGGTGATTGCCGTGGTGCTCTAGAA.

### Passaging of gDchi virus

Vero cells were infected with the HSV-1 gDchi viruses at a multiplicity of infection (MOI) of 0.01. After complete cytopathic effect (CPE) was observed, cells were harvested and sonicated to create a virus stock. Virus then was reseeded on Vero cells for serial passages in roller bottles. Large plaques were observed at passage 23.

### Microscopy and determination of plaque size

Plaques were visualized using Giemsa (Sigma-Aldrich, USA) staining after three days of infection and imaged with transmitted light microscopy using EVOS Cell Imaging Systems (AMG, Fisher Scientific) at 40X magnification. To measure plaque size, randomly selected plaques were enlarged on a screen and the average plaque radius was calculated from two independent measurements of at least 50 plaques. Using this average radius, the plaque area and ratio of plaque size between mutant and WT viruses was determined, as described by previous described by (25, 26).

### Cell-based ELISA (CELISA)

To evaluate the cell surface expression of gC mutants, CHO-K1 cells seeded in 96-well plates were transfected with 60 ng of empty vector or a gC construct using 0.15 µl of Lipofectamine 2000 (Invitrogen) diluted in Opti-MEM (Invitrogen). After 24 h, the cells were rinsed with phosphate buffered saline (PBS) and CELISA staining was performed as previously described (74), using the primary anti-HSV-1 gC MAb (EastCoast Bio, USA). After incubation with primary antibody, the cells were washed, fixed, and incubated with biotinylated goat anti-mouse IgG (Sigma), followed by streptavidin-HRP (GE Healthcare) and HRP substrate (BioFX).

### Cell-cell fusion assay

The fusion assay was performed as previously described (72). Briefly, CHO-K1 cells were seeded in 6-well plates overnight. One set of cells (effector cells) were transfected with 400 ng each of plasmids encoding T7 RNA polymerase, gB or a gB mutant, gD, gL, and either WT gH or a gH mutant, together with gC or a gC mutant, using 5 µl of Lipofectamine 2000 (Invitrogen, USA). A second set of cells (target cells) was transfected with 400 ng of a plasmid encoding the firefly luciferase gene under control of the T7 promoter and 1.5 µg of receptor (nectin-1), using 5 µL of Lipofectamine 2000. After overnight transfection, the cells were detached with versene and resuspended in 1.5 mL of F12 medium supplemented with 10% FBS. Effector and target cells were mixed in a 1:1 ratio and re-plated in 96-well plates for six hours. Luciferase activity was quantified using a luciferase reporter assay system (Promega) and a Wallac-Victor luminometer (Perkin Elmer) (Fig. 4 and 6).

### Complementation of gC-null viruses

The complementation assay was performed as previously described (43, 75–77). Briefly, Vero cells in 6**-**well plate were transfected with 2 µg of plasmids encoding WT gC, a gC mutant, or empty vector (pSG5) using lipofectamine 2000. After an overnight incubation at 37°C, cells were infected with a gC-null HSV-1 (41) at 5×10^6^ per well. After 2 hours at 37°C, the medium was removed and the extracellular virus was inactivated by a one-minute exposure to sodium citrate buffer at pH 3.0 (**24**). Fresh medium was added, and the cells were incubated at 37°C overnight. Plates were subjected to three freeze-thaw cycles, and cell lysates containing the complemented virus were titered on Vero cells (Fig. 5A).

### Entry of complemented gC-null viruses

To quantify virus entry (22, 66), M1A or M3A cells growing in 96-well plates were infected with complemented gC-null HSV-1 stocks (41) at an MOI of 1. After five hours at 37°C, the cells were washed with PBS and lysed in 50 μL/well DMEM containing 0.5% NP-40. β-galactosidase activity was measured by adding 50 μL/well of a 4.8 mg/mL solution of chlorophenol red-β-D-galactopyranoside (CPRG; Boehringer Mannheim) and reading the absorbance at 560 nm on a plate reader (Perkin Elmer). This assay was repeated at least three times (Fig. 5B).

### Statistical analysis

Statistical comparison of plaque areas was performed with a Mann-Whitney U test using SPSS (version 25). The rest of the statistical tests were two-tailed *t* tests using SPSS. Analyses were performed using IBM SPSS statistics version 25 for Windows (IBM Corp., Armonk, NY) and SAS 9.4 (SAS Institute, Cary, NC).

## ACKNOWLEDGMENTS

We thank Dr. Gregory Smith for providing the HSV-1 BAC GS3217, as well as Dr. Yasushi Kawaguchi for providing the parental BAC. We thank Nan Susmarski for excellent technical assistance and members of the Longnecker laboratory for their help in these studies. Sequencing services were performed at the Northwestern University Genomics Core Facility. R.L. is the Dan and Bertha Spear Research Professor in Microbiology-Immunology. Research reported in this publication was supported by the National Institute of Allergy and Infectious Disease (NIAID) of the National Institutes of Health under grant number AI148478. The content is solely the responsibility of the authors and does not necessarily represent the official views of the National Institutes of Health. This manuscript is the result of funding in whole by the National Institutes of Health (NIH). It is subject to the NIH Public Access Policy. Through acceptance of this federal funding, NIH has been given a right to make this manuscript publicly available in PubMed Central upon the Official Date of Publication, as defined by NIH.

## Data Availability Statement

All data and reagents are available upon request, please contact the corresponding author directly for reuse.

